# Association between Parkinson’s Disease subtypes and tests of physical function: the 360-degree turn test is most predictive

**DOI:** 10.1101/342733

**Authors:** Morgane Prime, J. Lucas McKay, Allison Bay, Ariel Hart, Chaejin Kim, Amit Abraham, Madeleine E. Hackney

**Affiliations:** Faculty of Biology, University of Toulouse III - Paul Sabatier, Toulouse, France; Department of Biomedical Engineering, Emory University and the Georgia Institute of Technology, Atlanta, Georgia, USA; Division of General Medicine and Geriatrics, Department of Medicine, Emory School of Medicine, Atlanta, Georgia, USA; Department of Kinesiology, University of Georgia, Athens, GA, USA; Atlanta VA Center for Visual and Neurocognitive Rehabilitation, Decatur, Georgia, USA; Department of Rehabilitation Medicine, Emory School of Medicine, Atlanta, Georgia, USA

**Keywords:** Parkinson’s disease, motor subtypes, postural instability and gait difficulty, tremor dominant, 360-degree turn test

## Abstract

**BACKGROUND AND PURPOSE:** People with Parkinson’s disease (PD) present phenotypes that can be characterized as tremor-dominant (TD) or postural instability / gait difficulty (PIGD) subtypes. Differentiation of subtypes allows clinicians to predict the disease course and adjust treatment accordingly. We examined whether brief mobility and balance measures can discriminate PIGD from TD phenotypes.

**METHODS:** We performed a cross-sectional study with individuals with PD (N=104). Blinded raters assessed participants with the UPDRS or MDS-UPDRS, and potential predictor variables: 360-degree turn test, one-leg stance, backward perturbation test and tandem walk. Participant were classified as PIGD or TD based on the Unified Parkinson’s Disease Rating Scale or the Movement Disorder Society revision (UPDRS or MDS-UPDRS) assessment results. Differences in study variables between subtype groups were assessed with univariate analyses. Receiver operating characteristic (ROC) curve analyses were performed to investigate the ability of candidate predictor variables to differentiate PD subtypes.

**RESULTS:** Mean age and disease duration were 68±9 and 7±5 years, respectively, and Hoehn & Yahr Stages I-IV median (1st,3rd quartile) = II (II, III). No differences between subtypes were observed for tandem walk or reactive postural control. PIGD participants performed worse on number of steps (p<0.001) and time to complete (p=0.003) the 360-degree turn test and one-leg stance (p=0.006). ROC curves showed only the 360-degree turn test could discriminate PIGD from TD with high sensitivity.

**CONCLUSIONS:** The 360-degree turn test requires minimal time to administer and may be useful in mild-moderate PD for distinguishing PIGD from TD subtypes.

## INTRODUCTION

People with Parkinson’s Disease (PD) experience bradykinesia, rigidity, resting tremor, postural instability and balance perturbation. Such motor impairments lead to functional disabilities,^1^ including high risk of falls^1^ which increases with disease severity and often burdens patients and caregivers.

PD is quite heterogeneous in terms of age of onset, side dominance and clinical manifestations. Compared to early-onset PD persons, late-onset PD persons more often exhibit postural instability and gait difficulty (PIGD) symptoms^2^ and more impaired cognition.^3^ In addition, symptoms like bradykinesia and PIGD are more common in persons with more rapid disease progression than those with a slower disease progression.^3^

PIGD phenotype is mostly characterized by bradykinesia and rigidity in movement^4^ whereas tremor-dominant (TD) persons typically display resting tremor, normal gait, and mild disease progression.^5^ Comparisons of TD with PIGD-dominant PD phenotypes supported the existence of clinical subtypes,^3^ each with distinct motor features and prognoses.^6^ PIGD persons experience significantly greater subjective intellectual, motor and occupational impairment than TD persons^3^ and have a higher incidence of dementia^7^ and neuropsychiatric disorders such as depression^8^ and apathy.^9^ Differentiation between TD and PIGD parkinsonism facilitates a better understanding of their deficits and impacts, allowing clinicians to tailor interventional strategies that accommodate patients’ unique symptomatology and changing needs.^10,11^ For example, although the responsiveness to both subthalamic nucleus (STN) and globus pallidus internus (GPi) deep brain stimulation (DBS) is similar among the two motor subtypes, TD persons may exhibit a greater response to GPi DBS with respect to gait and PIGD persons display less overall benefit from stimulation.^12^

PD motor deficits, especially postural instability and gait difficulties, commonly lead to poor performance in measures of balance and functional tasks, and have been associated with falls.^13^ The Unified Parkinson’s Disease Rating Scale or the Movement Disorder Society revision (UPDRS or MDS-UPDRS) are used to distinguish TD persons from PIGD persons based on assessment scores.^14,15^

Algorithms and formulas have been developed to define TD and PIGD phenotype using their UPDRS and MDS-UPDRS.^3^ The ratio of the mean UPDRS tremor scores to the mean UPDRS PIGD scores are used to define TD persons (ratio 1.5) and PIGD persons (ratio 1.0), with a group of persons in between these scores that is referred to as “indeterminate.” This common classification method has been demonstrated to be a reliable way to classify persons by motor subtype.^14^ However, UPDRS and MDS-UPDRS take a long time to administer.

Common scales assessing balance -- Fullerton Advanced Balance (FAB) scale, Mini-Balance Evaluation Systems Test (Mini-BESTest) and Berg Balance Scale (BBS) -- moderately distinguish fallers from non-fallers in individuals with PD. Schlenstedt et al. found that specific items of these three balance scales were useful in predicting future falls. They recommended using a model combining “tandem stance,” “one-leg stance,” “rise to toes,” “compensatory stepping backward,” “turning 360-degree” and “placing foot on stool” when analyzing postural control deficits. ^16^ This recommendation indicates that these specific functional tasks may have predictive power that can shed light on postural control abilities. Studies comparing postural control in non-fallers versus fallers have shown that persons with PD who fall have reduced reactive postural control^17–19^ and an impaired ability to perform tandem stance/walk.^20^ Fallers are reported to present poorer ability to stand on one leg.^21^ Finally, an increase in the number of steps and time taken to complete a 360-degree turn is suggested to be a compensatory strategy that people use to avoid loss of balance and prevent falls.^22^

Such functional measures are advantageous because they can be quickly administered in clinical and research settings, particularly when a comprehensive neurological examination is infeasible or when item-level motor examination results are unavailable. Therefore, it may be appropriate to assess PD persons with similar balance assessments and examine differences between PIGD and TD participants. Receiver operating characteristic (ROC) analysis is an approach for examining the sensitivity and specificity of each predictor variable to differentiate between subgroups.^23^ Having short and simple tests to identify persons’ phenotypes with high sensitivity and specificity would enable clinicians to efficiently determine intervention plans.

This cross-sectional study was performed using baseline measurements of N=104 PD participants who had volunteered for rehabilitation or observational research studies. The first aim was to compare performances of each motor subtype (i.e. PIGD and TD) to physical tests derived from common balance measures. The second aim was to examine whether these brief physical examinations could predict and differentiate TD and PIGD phenotypes. Participants were classified according to motor subtype with algorithms using the UPDRS or the MDS-UPDRS.^14^ Quantitative measures collected as potential predictor variables included: time and number of steps needed to complete a 360-degree turn, the amount of time a participant could stand on one leg, the number of interruptions in tandem walk and number of steps taken during reactive postural control. ROC curve analyses were implemented to assess the ability of potential behaviorally-derived predictor variables to differentiate PD subtypes. As PIGD subtype is associated with more functional disability than TD subtype,^3^ PIGD persons were expected to have worse performance i.e. taking more time and more steps to complete a 360-degree turn, standing less time on one-leg, execute tandem walk with more interruptions and more steps to recover their balance during reactive postural control.

## METHODS

### Study

This study was approved by the Institutional Review Board at Emory University School of Medicine and the Research & Development Committee of the Atlanta Veterans Affairs (VA). Participants provided written informed consent before participating.

### Participants

Participants were recruited through the VA registry, the VA Informatics and Computing Infrastructure (VINCI) database, the Michael J. Fox FoxFinder website, the Movement Disorders unit of Emory University, PD organizations’ newsletters, support groups and educational events at the local community and through word of mouth. Participants had a clinical diagnosis of PD made by a movement disorders specialist based on the United Kingdom PD Society Brain Bank diagnostic criteria. ^24^ They exhibited three out of the four PD cardinal signs (postural instability, tremor, rigidity and bradykinesia), had unilateral onset of symptoms and had shown clear symptomatic benefit from antiparkinsonian medications, e.g., levodopa.^25^ Participants were aged 40 and older, were stage I-IV in the Hoehn & Yahr scale and could walk three meters or more with or without assistance.

### Study variables

Each participant was assessed with a standardized battery that included measurements of disease severity, global cognition^26^^,27^ and physical function (Composite Physical Function Index, “CPF”).^28^ Testing was performed in the “OFF” state, i.e., at least 12 hours after their last antiparkinsonian medication to reduce the impact of medication-related motor fluctuations.

The methods suggested by Stebbins et al. were employed to classify participants as either PIGD or TD. Among those participants assessed with the MDS-UPDRS (n=74), a tremor subscore was assembled as the mean value of items II.10, III.14 LUE, III.15 RUE, III.16 LUE and RUE. A PIGD subscore was assembled as the mean value of left upper extremity (LUE), right upper extremity (RUE), left lower extremity (LLE), and right lower extremity (RLE) items II.12, II.13, III.10, III.11 and III.12. Among those participants assessed with the UPDRS (n=30), the tremor subscore was assembled as the mean value of items II.16, III.20 face, RUE, LUE, RLE and LLE, III.21 RUE and LUE. and the PIGD subscore was assembled as the mean value of items II.13, II.14, II.15, III.29 and III.30. The ratio of the tremor to PIGD subscores was then used to classify participants as TD (>1.15), PIGD (<0.90), or otherwise indeterminate. Total UPDRS-III scores were transformed into MDS UPDRS-III scores prior to univariate analyses according to methods in the literature. ^15^

Candidate predictor variables were derived from specific items of the FAB assessment tool: the 360-degree turn test, tandem walk, one-leg stand, and reactive postural control. For the 360-degree turn test, subjects were asked to turn around in a full circle, pause and then turn in a second full circle in the opposite direction. During the tandem walk, participants had to walk forward along a line, placing one foot directly in front of the other such that the heel and toe were in contact on each step forward. Participants were also asked to fold their arms across their chest, lift a leg off the floor without touching the other leg and stand with eyes open as long as they could. To test their reactive postural control, participants slowly leaned back into rater’s hand until the rater removed their hand and counted the number of steps the subject needed to recover their balance.

Missing data were as follows. Among PIGD participants: sex and age (n=1), education (n=2), ethnicity (n=1), housing and transportation (n=2), medication (n=1), number of years with PD (n=1), Use of assistive device for walking (n=2), self-rated quality of life (n=2), frequency of leaving house (n=3), composite physical function score (n=3), freezing more than once a week (n=5), MDS-UPDRS Part I, II and IV (n=21). Among TD participants: MDS-UPDRS Part I, II and IV (n=9).

### Statistical analysis

All available data were used for participants, including those for whom data for some assessments are unavailable.

First, data were inspected for normality with SPSS statistical software (IBM SPSS Statistics version 24). The 360-degree turn test steps, tandem walk, one-leg stance, reactive postural control, and MoCA all had skewness and kurtosis statistics less than the widely acceptable +/-2, showing a normally distributed univariate distribution.^29^ The skewness and kurtosis statistics for the 360-degree turn test time did not follow a normal distribution; however, upon further exploration it was determined that this was due to the presence of outliers (where the 360-degree turn test took more than 10 seconds to complete, n=9). Once the outliers were removed, skewness and kurtosis were 1.00 and 0.56, respectively. For many participants, the long turn time was due to freezing of gait. For example, one participant experiencing freezing took more than 45 seconds to complete the 360-degree turn. Although the outliers caused data to be skewed, freezing of gait is a common symptom of PD, especially among the PIGD phenotype; therefore, the outliers were included in the ROC analysis because exclusion of the outliers would limit external validity.

Participants who were classified as ‘indeterminate’ (n=12) were excluded from further analyses in order to be able to detect clear differences between ‘TD’ and ‘PIGD’ group. Separate univariate logistic regression analyses were applied to identify associations between PD phenotype (PIGD vs. TD, with TD as the reference group) and each of the following candidate predictor variables: Number of Steps to Turn 360-degree, Time to Turn 360-degree (s), Number of Interruptions in Tandem Walk, Maximal Time Standing on One Leg (s), Number of Steps to recover balance during Reactive Postural Control. Identified logistic regression models were used to create receiver operating characteristic (ROC) curves for each candidate predictor variable plotting the sensitivity vs. 1-specificity of the model to predict membership in the PIGD group.^23^ Furthermore, optimal cutoff points were investigated maximizing both sensitivity and specificity. Chosen type-I error was alpha = 0.05. Rstudio software (version 1.0.153) was used for the statistical analysis.

## RESULTS

104 PD persons with either TD or PIGD phenotype were included in this study. Table 1 shows the characteristics of the sample. Males represented 58.3% of the sample (n=60), which is representative of PD epidemiology.^30^ Mean age and disease duration were 68±9 and 7±5 years, respectively. Most participants were between stage 1.5 and stage 3 on the Hoehn & Yahr scale. PIGD participants significantly reported more freezing episodes per week and more falls in the previous six months than TD participants. They also used more assistive devices for walking. Compared to PIGD participants, TD participants had better composite physical function scores and MoCA scores, reflected by a higher self-reported quality of life.

**Table 1.**
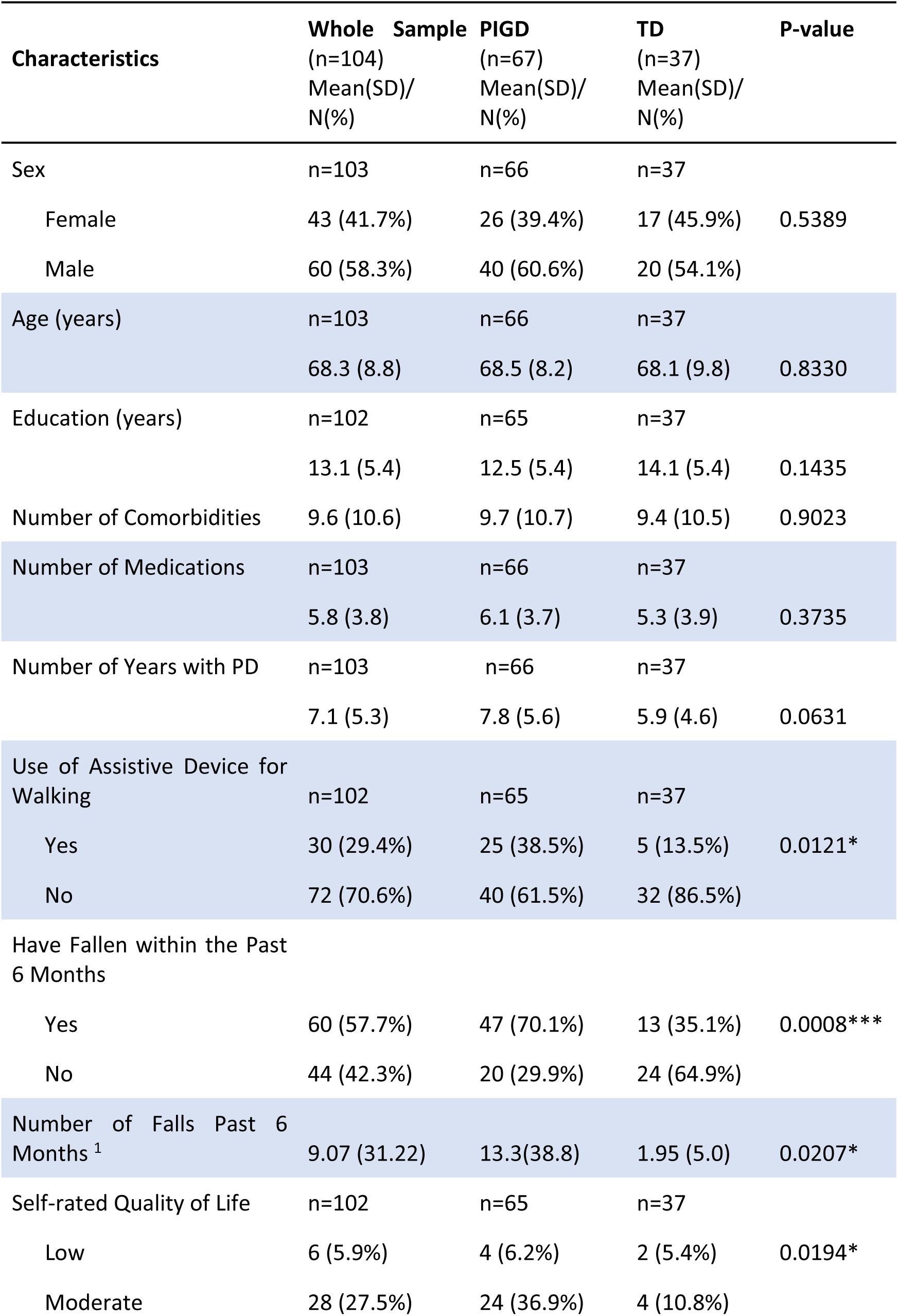
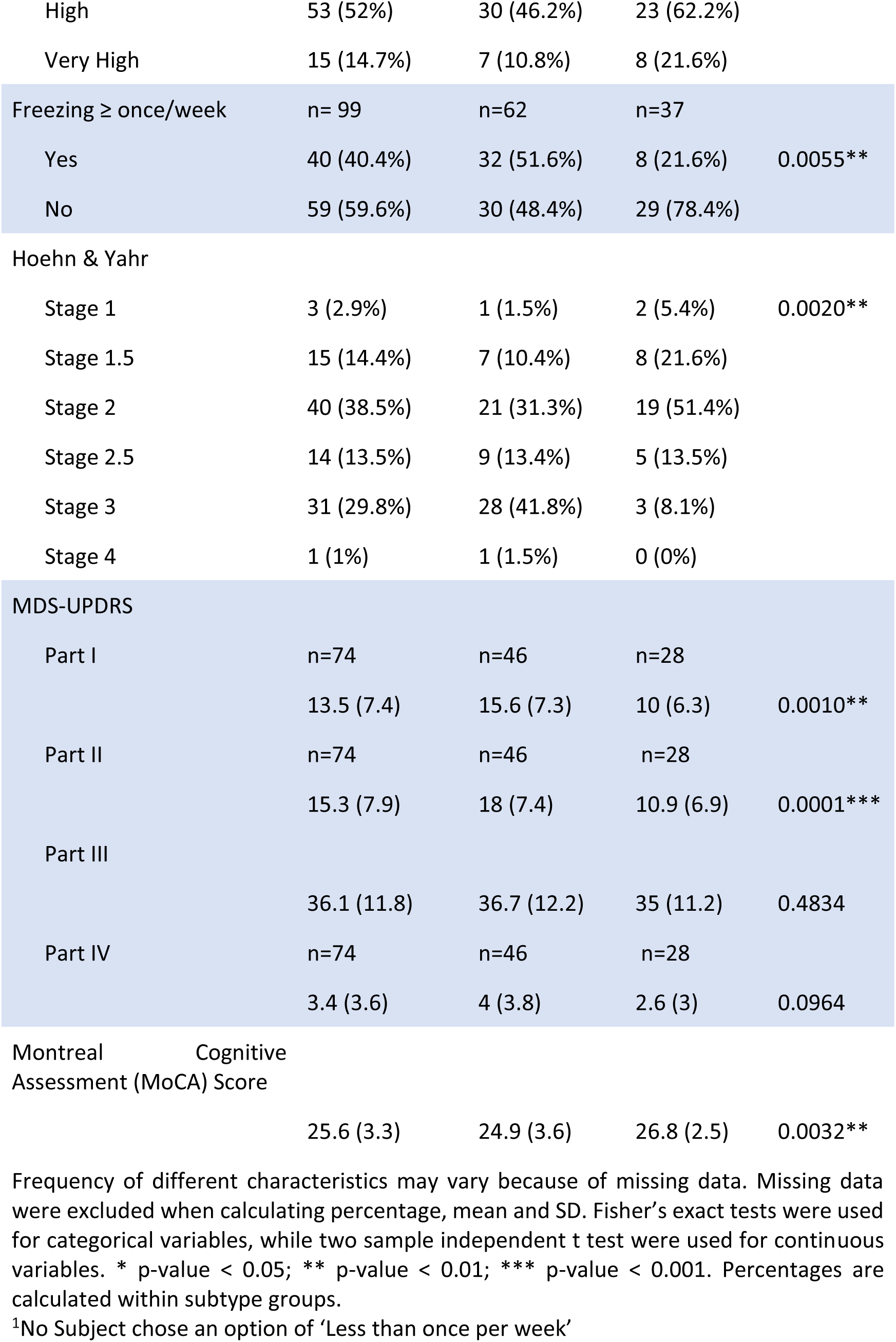
Demographic Characteristics of Parkinson’s Disease Motor Subgroups (Tremor Dominant and Postural Instability/Gait Difficulty).

Univariate associations between individuals’ motor subtype and physical measures are provided in table 2 and figure 1. This analysis provides evidence that PIGD participants needed more time (p=0.003) and more steps (p<0.001) to complete the 360-degree turn test. PIGD participants had shorter one-leg stance times (p=0.006). Number of steps taken in reactive postural control and number of interruptions during the tandem walk did not differ between groups.

**Table 2.**
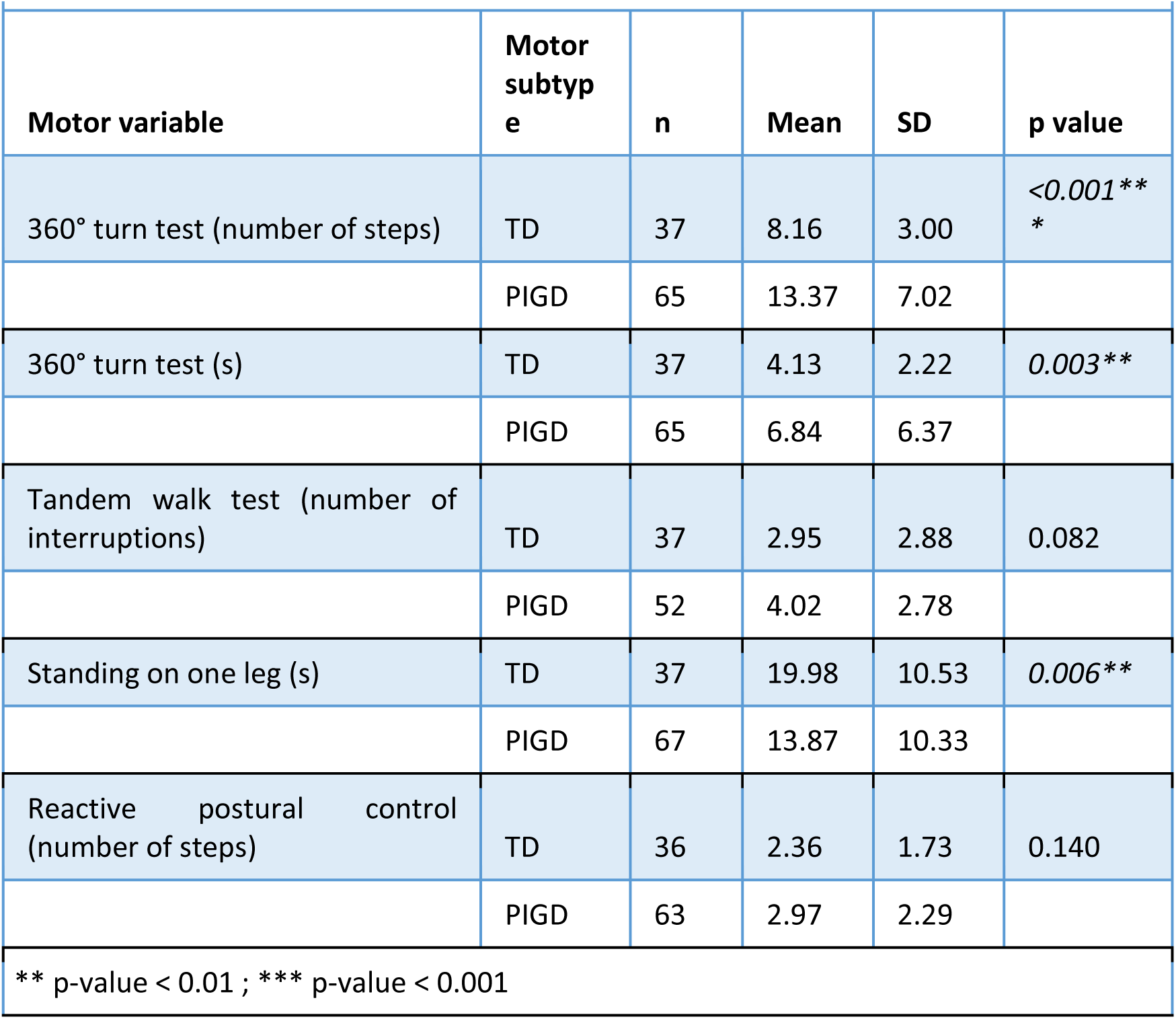
Tremor Dominant and Postural Instability/Gait Difficulty participants’ performance on Physical Exams.

**Figure 1.**
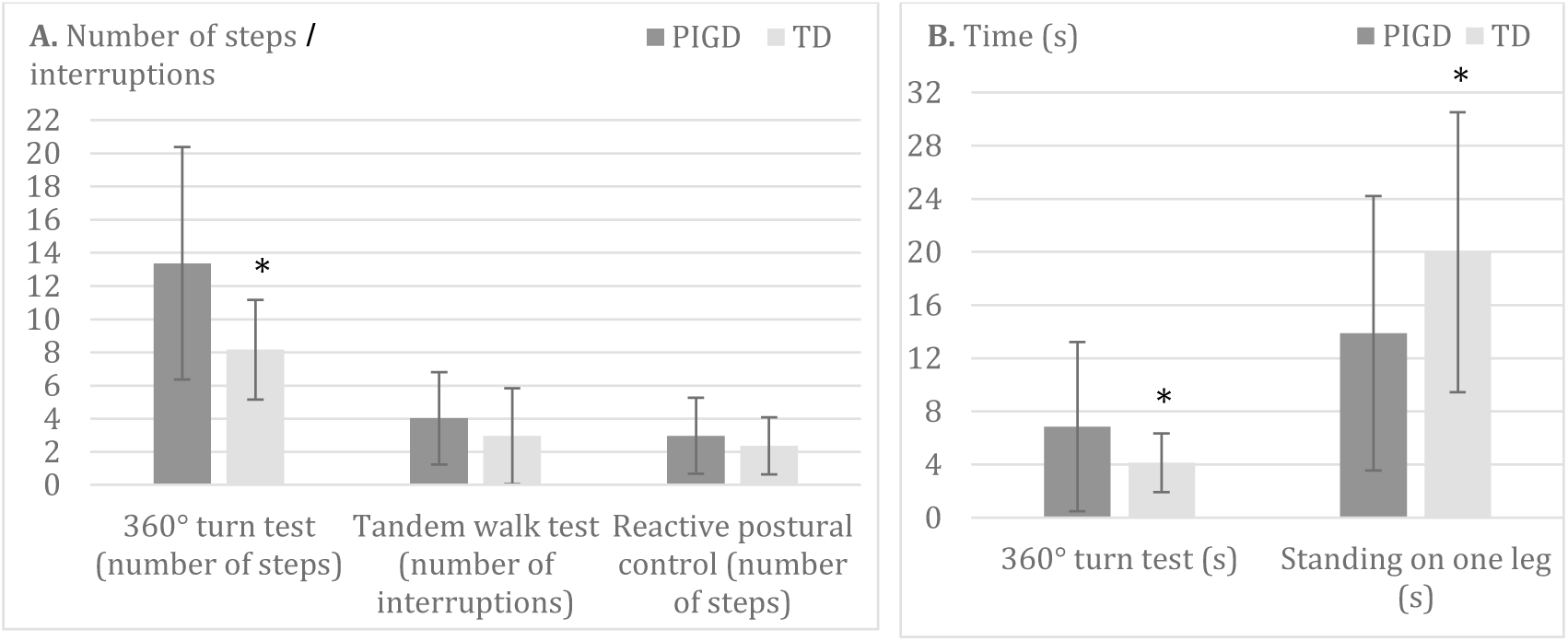
Tremor Dominant and Postural Instability/Gait Difficulty participants’ performance on Physical Exams. **A.** Number of Steps to Turn 360-degree, Number of Interruptions in Tandem Walk, Number of Steps to recover balance during Reactive Postural Control. **B.** Time to Turn 360-degrees (s), Maximal Time Standing on One Leg (s). *p-value <0.05. Values plotted are means ± standard deviation.

ROC curves are shown in figure 2 with results of their analysis including: AUC, confidence intervals (CI), p-value, sensitivity, specificity and optimal cut-off point for each test are displayed in table 3. As individuals with indeterminate phenotypes were excluded from analysis, participants who were not identified as TD phenotype are PIGD phenotype. “Optimum cutoff” refers to the threshold for classification between TD and PIGD such that values greater than the optimum cutoffs are classified as PIGD. With an AUC of 0.580, reactive postural control does not enable discrimination of PIGD from TD. According to the p-value of 0.007 for the ROC curve analysis, the one-leg stance can predict participants’ motor subtype. However, AUC of 0.654 and specificity of 0.676 are quite low. Only one test is outstanding in this ROC curve analysis: the 360-degree turn test. Using number of steps and time to achieve a complete turn is a way to identify PIGD persons with an accuracy of respectively 0.751 and 0.740 for the AUC and 0.838 and 0.892 for the sensitivity.

**Table 3.**
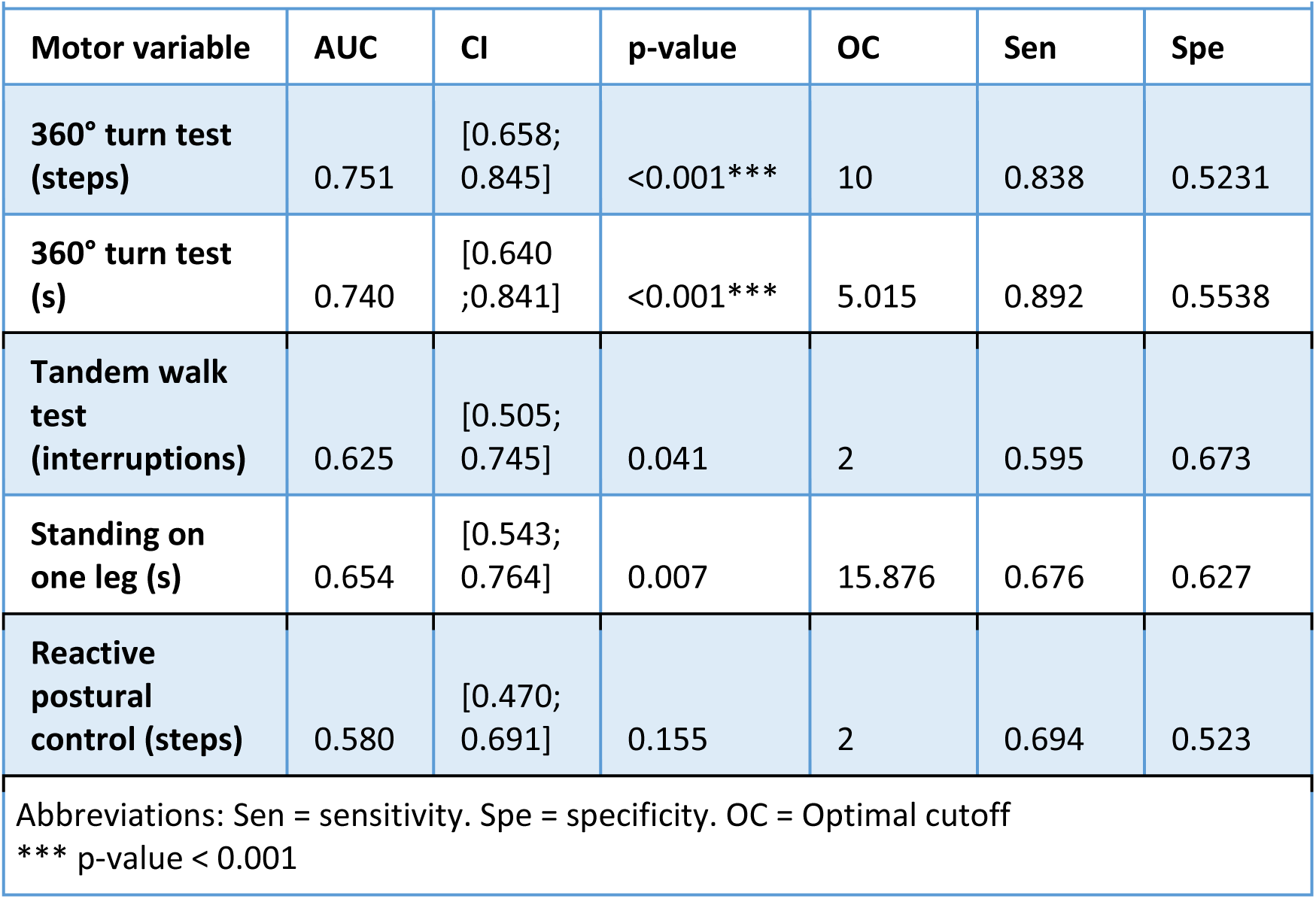
Receiver Operating Characteristics curve analysis of Simply Physical Exams for TD phenotype.

**Figure 2.**
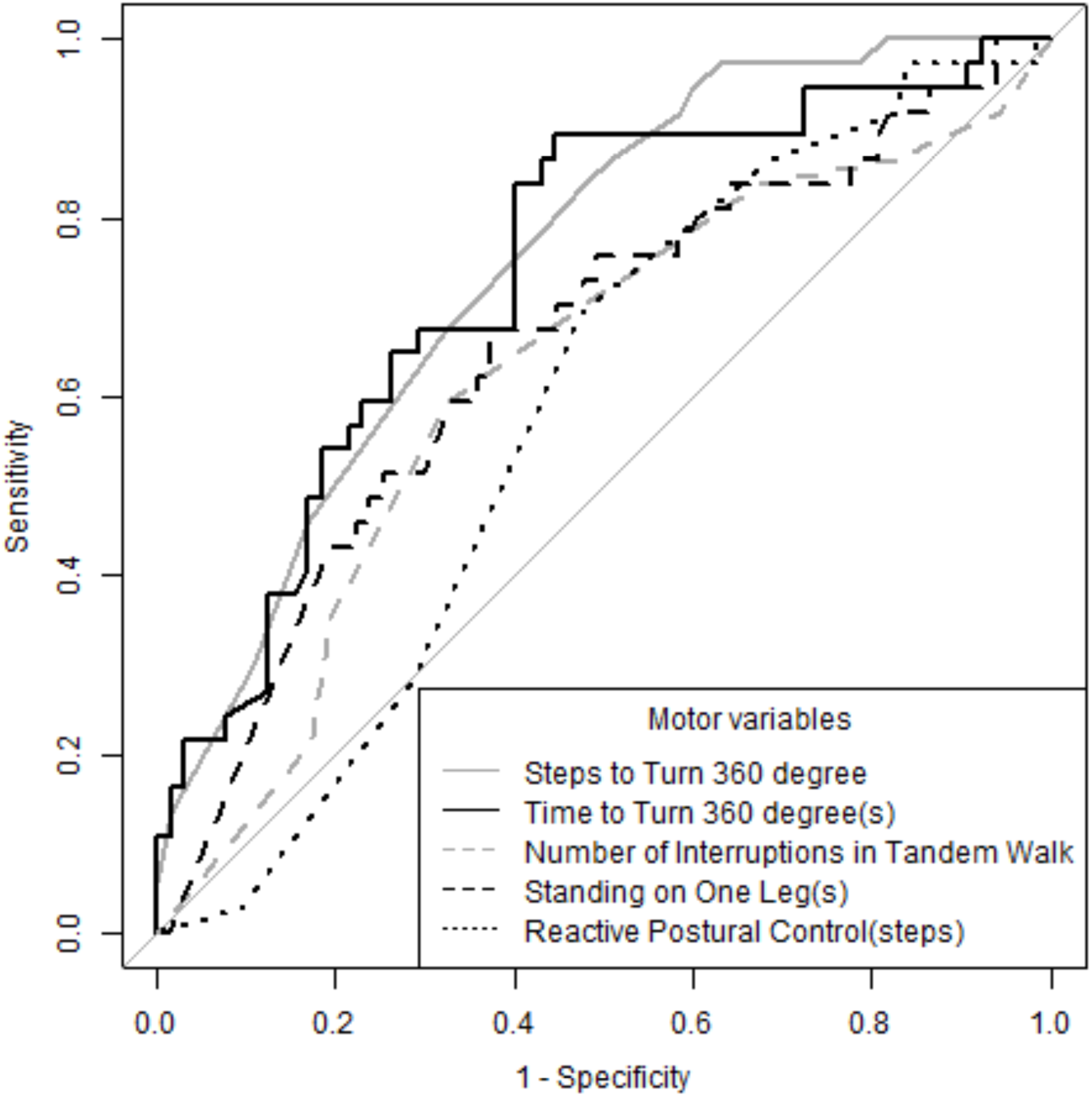
Receiver Operating Characteristic Curve of Physical Exams for TD Phenotype.

This analysis suggests that steps and time to turn 360-degree can correctly identify a person with PD as the PIGD phenotype in 83.8% and 89.2% of cases examined, respectively. In addition to the 360-degree turn test, the tandem walk test and one-leg stance had significant “area under the curve” (AUC) values, which suggests that these tests are significant predictive variables; however, only the 360-degree turn test is sufficiently sensitive (>0.80) to be clinically useful.

## DISCUSSION

This study revealed that among several standard, clinical and validated tests of postural stability and mobility, only the 360-degree turn test was able to distinguish PIGD subtype from TD subtype with high sensitivity using both number of steps and time. Thus, this test could serve as a quick and easily administered screening tool for clinicians to identify people with PIGD, who arguably have a more severe prognosis. However, the 360-degree turn test has low specificity, which implies that some patients who are actually TD would be labeled as PIGD. Because of the trade-off between specificity and sensitivity - as the ideal situation of a 100% accurate test is unrealistic, Lalkhen and McCluskey recommend assessing people who are initially positive in such situations with an additional test with low sensitivity and high specificity.^31^ Further investigation is needed to find another short and simple physical test that could discriminate PIGD from TD phenotype with high specificity.

According to the current results, reactive postural control cannot be a predictor of PD motor subtype as TD and PIGD phenotype performed similarly on this test. Although tandem walk seems to be predictive for PIGD subtype with a p-value of 0.041 for the ROC curve in the present sample, the t-test showed no significant differences in number of interruptions between TD and PIGD phenotypes. Furthermore, sensitivity in ROC curve analysis of tandem walk is relatively low (0.595). This suggests that the number of interruptions during a tandem walk should not be considered as a tool to discriminate PIGD persons.

Previous studies investigating the relationship between postural control and falls also showed conflicting results. Dennison et al. validated tandem walking as a predictor of recurrent falls (p<0.006) whereas de Oliveira Souza et al. demonstrated that there was no significant correlation between tandem walk and tests assessing clinical balance and executive function.^32,33^ One-leg stance was moderately able to distinguish PIGD from TD (sensitivity=0.676). This physical test is mostly a predictor of fallers. However, PIGD persons tend to fall more than TD persons, reflecting greater balance impairments, which can explain the correlation between motor subtype and one-leg stance performances in their study.

Turning is impaired in people with PD, even in mildly affected individuals. Stack and Ashburn showed that PD persons need a mean of seven steps to turn. ^34^ Compared to the five steps normally required by elderly populations, an increase in the number of steps has been correlated to UPDRS and Self-assessed Disability Scale (SAS) scores, demonstrating the impact of disease severity.^35^ As well as significantly impacting quality of life, turning difficulties have been related to poor cognitive functioning^36^ and are a sensitive predictor of the two key symptoms of PD locomotion: freezing - a sudden interruption of ongoing movement^35^ and falling.^37^ Freezing of gait (FOG) and falls being predominant in PIDG persons, this study showed that the 360-degree turn test may be a quick and easy way to distinguish PIGD from TD. Although increased number of steps and time to complete a 360-degree turn reflect functional impairments, this strategy is hypothesized to be compensatory on the part of the participant to maintain their balance and avoid falls during the turn.^22^

The motor tests explored in the current study included a combination of dynamic balance and static balance activities, which serve as possible predictive variables for subtypes. These different balance tasks activate distinct regions of the brain. Studies have described the involvement of the sensorimotor cortex (SMC) in normal gait, ^38,39^ and the supplementary motor area (SMA) and prefrontal cortex (PFC) for walking speed.^40^ One possible explanation for the ability of the 360-degree turn test and the one-leg stance to differentiate between PD phenotypes may be that these tests capture the functional impacts of the different patterns of neurodegeneration between phenotypes. One-leg stance appears to engage activity in the parietal cortex.^41^ Furthermore, TD and PIGD phenotypes have differing patterns of neurodegeneration. At same disease stages and duration, PIGD persons exhibit more white matter degradation, which supports the idea of TD being a less pathological PD subtype.^42^ Given that the 360-degree turn test provided the most sensitive discriminatory ability, it is possible that the differing patterns of neurodegeneration have greater functional impacts on dynamic than static balance activities. As such, these findings underscore the importance of evaluating dynamic balance activities in addition or even instead of static balance activities.

The findings of this study have clinical implications. Differing therapeutic interventions are recommended for TD and PIGD phenotypes to address the phenotype-specific challenges associated with PD.^43^ Although PD phenotype is commonly derived from the UPDRS and MDSUPDRS, these tools contain many questions and require a long period of time to complete. Clinicians and researchers who are interested in tailoring interventions to the individual would likely benefit from having shorter assessments to determine PD phenotype. Conducting the 360-degree turn test may provide a simple but effective way to differentiate between subtypes as opposed to conducting the entire MDS-UPDRS/UPDRS assessment, allowing clinicians and PD patients to conserve time while expediting the development of individual rehabilitation regimens.

## LIMITATIONS

The 360-degree turn test was administered in real time and a trained rater counted the steps and timed the turns with a stopwatch. Some have said that capturing this information with inertial sensors would lead to more accurate estimations of performance.^44^ However, it has also been demonstrated that timing mobility measures, e.g., gait speed, can be accurately and reliably timed with a stopwatch. ^45^

A few values for multiple variables are missing for several participants. They either did not want to give the information or more frequently they inadvertently skipped answering some items in the questionnaires and these mistakes were not caught by examiners. Regarding the physical examination, participants sometimes refused to perform some tests for various reasons, i.e. they felt they could not attempt the tests given their level of mobility. Notably, participants often fear falls. It is difficult to know whether persons performed poorly because of their lack of trust in their balance or because they truly have poor balance mechanisms. Mak and Pang showed that balance confidence and functional mobility are independently associated with falls in people with PD.^46^ These missing data are limitations; however, we did not use imputations, in order not to mislead interpretations in one way or the other.

The internal validity may be compromised because our sample size was not big enough to split the data into training and testing sets. External validity is also limited because of the exclusion of indeterminate PD persons. Furthermore, ROC analysis used to determine the optimal cut-off points has been suggested to limit the generalizability of results.^47,48^

## CONCLUSION

The 360-degree turn test requires minimal time to administer and may be useful in mild-moderate PD for distinguishing PIGD from TD subtypes, particularly when comprehensive neurological examination is infeasible or when item-level motor exam results are unavailable.

## Acknowledgement

We would like to thank the participants for taking part in this study as well as research assistants, medical students and undergraduate volunteers at Emory School of Medicine for data collection, entry and verification.

## SUPPLEMENTAL INFORMATION

**Table 4.**
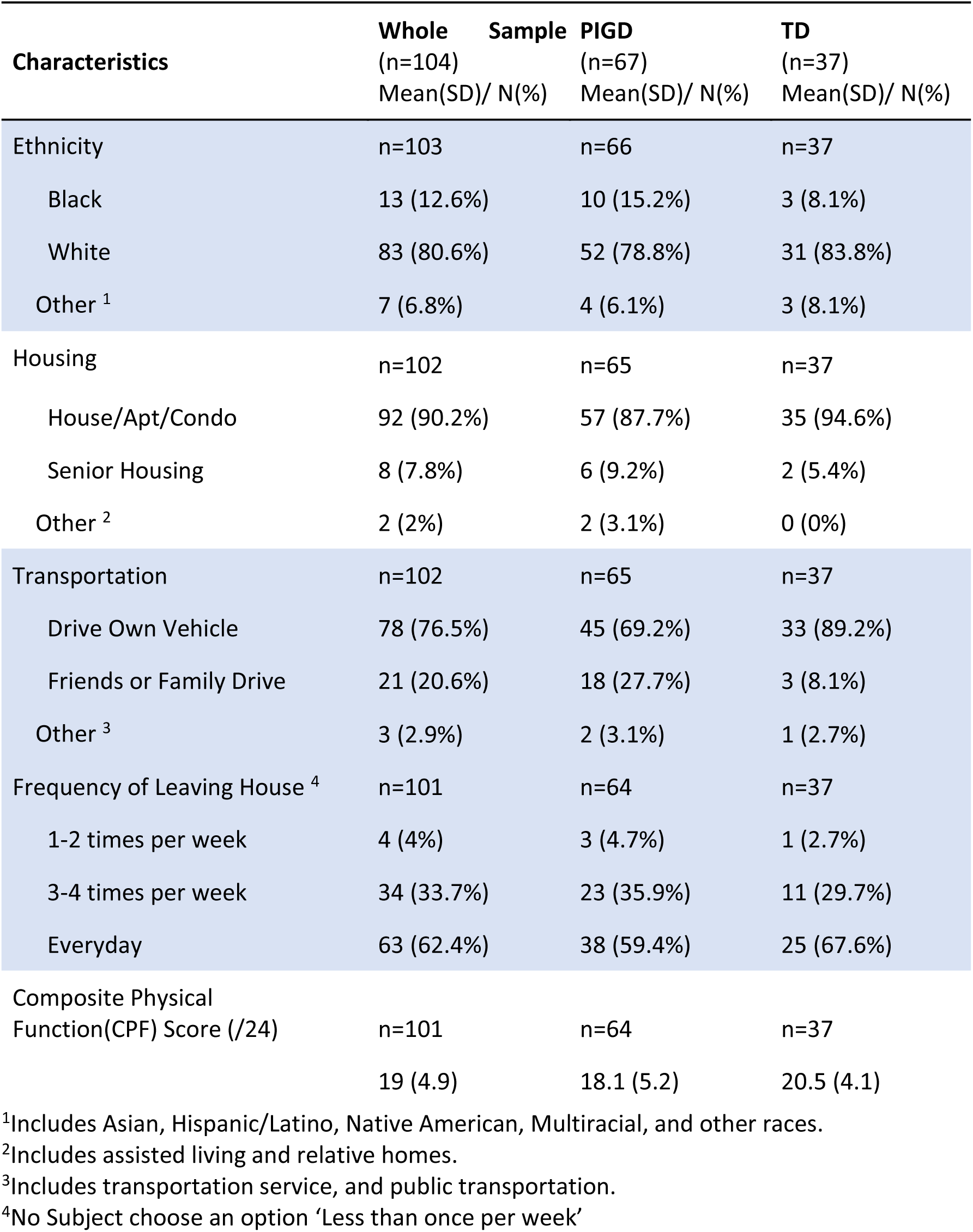
Other Demographic Characteristics by Parkinson’s Disease Motor Subgroups (Tremor Dominant and Postural Instability/Gait Difficulty).

